# Postsynaptic Pannexin-1 Facilitates Anandamide Uptake to Modulate Glutamate Release and Enhance Network Excitability

**DOI:** 10.1101/263640

**Authors:** Jennifer Bialecki, Nicholas L. Weilinger, Alexander W. Lohman, Haley A. Vecchiarelli, Jordan H.B. Robinson, Jon Egaña, Juan Medizabal-Zubiaga, Allison C. Werner, Pedro Grandes, G. Campbell Teskey, Matthew N. Hill, Roger J. Thompson

## Abstract

Prolonged neurotransmitter release following synaptic stimulation extends the time window for postsynaptic neurons to respond to presynaptic activity. This can enhance excitability and increase synchrony of outputs, but the prevalence of this at normally highly synchronous synapses is unclear. We show that the postsynaptic channel, pannexin-1 (Panx1) regulates prolonged glutamate release onto CA1 neurons. Block of postsynaptic (CA1 neuronal) Panx1 increased the frequency of glutamate neurotransmission and action potentials in these neurons following Schaffer collateral stimulation. When Panx1 was blocked, anandamide levels increase and activated transient receptor potential vanilloid 1 (TRPV1)-mediated glutamate release. This TRPV1-induced synaptic acitvity enhanced excitability and translated into a faster rate of TRPV1-dependent epileptogenesis induced by kindling. We conclude that Panx1 facilitates AEA clearance to maintain synchronous release onto CA1 neurons so that when AEA clearance is reduced, TRPV1 channels prolong glutamate neurotransmission to enhance network output to promote epileptiform activity.

---

Fast synaptic transmission, in contrast to stimulation-independent ongoing spontaneous transmission, is characterized by the highly synchronous release of neurotransmitter in response to presynaptic activation ^1^. Asynchronous neurotransmitter release while less well understood, is typically stimulation-dependent and lasts hundreds of milliseconds. At excitatory synapses, asynchronous glutamate release can elevate firing rates in response to presynaptic stimulation ^2,3^, enhancing coincidence detection ^4^, and possibly promote spread of neurotransmitter ^1^. Aberrant asynchronous release may influence neurodegeneration ^5^ or promote epileptogenesis ^6,7^. At GABAergic synapses asynchronous release is likely involved in periods of prolonged inhibition ^8–10^. All synapses show spontaneous release and most are specialized for either synchronous or asynchronous release, although some switch from being predominantly asynchronous to synchronous during development ^1,4,11^. The demonstration that a typically synchronous synapse can switch between predominant modes of release would greatly expand their signalling capacity during physiology and pathology.

Asynchronous release typically involves a presynaptic Ca^2+^ permeable channel that is likely distal from the release machinery, resulting in more prolonged intracellular Ca^2+^ rises compared with voltage dependent Ca^2+^ channel activity and synchronous release ^1,11^. A common thread for these presynaptic Ca^2+^ sources is that they are regulated by extracellular ligands, such as ATP activation of P2X2 at CA1-interneuron synapses ^12^ and endovanniloids for TRPV1 afferents of the nucleus tractus solitaris (NTS) ^13,14^. The prevailing view is that these messengers function as either anterograde (presynaptic source), retrograde (postsynaptic source), or glial sources and it follows that the endocannabinoids/endovanilloids must be removed rapidly from the synapse for signal termination.

Pannexin-1 (Panx1) are ion / metabolite channels with well characterized pathological roles that include neuronal death during ischemia ^15–18^ and inflammation ^19^. Panx1 expression is reported in the postsynaptic density (PSD) of hippocampal and cortical pyramidal neurons ^20^, but the breadth of roles of these channels remain poorly understood. Panx1 enhances high frequency stimulation induced long-term potentiation (LTP) in the hippocampus ^21^ and attenuates low frequency induced long-term depression (LTD) ^22^. Panx1 contributes to aberrant excitability by increasing the amplitude and frequency of interictal bursts ^23,24^. Several acute, chemically induced seizure models have provided conflicting results on the ability of Panx1 to promote or inhibit ictal events ^25–27^. Single, acute chemically-induced seizures (i.e. via pilocarpine or kainic acid) provide information on the short-term role of Panx1. These models however provide little mechanistic insight into the progressive increase in seizure severity seen during epileptogenesis, which may involve asynchronous neurotransmitter release ^6,7,28^.

Here we report that block of postsynaptic Panx1 paired with Schaffer collateral simulation prolongs glutamate neurotransmission onto CA1 neurons that depends upon activation of presynaptic TRPV1 channels by the signalling lipid, anandamide (AEA). Panx1 regulates tissue levels of AEA by facilitating its clearance. Prolonged release increased tissue excitability and was critical for epileptogenesis during electrical kindling. Thus, we have found a novel form of short-term plasticity requiring postsynaptic Panx1 and presynaptic TRPV1 that shifts the normally highly synchronous glutamate release onto CA1 neurons into a prolonged mode and promotes pathological network synchrony.

## Results

### Block of postsynaptic Panx1 causes prolonged evoked glutamate release

Panx1 is expressed in the postsynaptic density ^20^ and may therefore regulate synaptic activity. We selectively inhibited Panx1 in single postsynaptic CA1 neurons by inclusion of a blocking antibody, αPanx1 (0.25 ng/μl) in the patch pipette ^17,18^. The control was a polyclonal antibody against connexin-43 (αCx43), which is not expressed in these cells, but like Panx1 is a member of the gap junction family ^18^. Postsynaptic block of Panx1 did not change basal synaptic activity because the frequency and amplitude of spontaneous excitatory postsynaptic currents (sEPSC) were unaltered (Fig. 1ab). Schaffer collateral stimulation (paired-pulse stimulations at 0.33 Hz, 1 ms duration, 50 ms interstim interval) induced a surprising increase in the frequency, but not the amplitude of sEPSCs, suggesting a presynaptic origin (Fig 1ab; note that shaded regions in b represent the stim population; longer exemplar recordings are shown in Fig. S1). These prolonged synaptic events occurred in all CA1 neurons tested (n=28). Interestingly, the prolonged synaptic events did not occur after every stimulation but where seen 22% (89/420) of the time and lasted for 13±0.5 s once initiated. Substituting αPanx1 for αCx43 (0.3ng/μl) did not induce prolonged release upon Schaffer collateral stimulation (Fig. 1b).

**Fig. 1.**
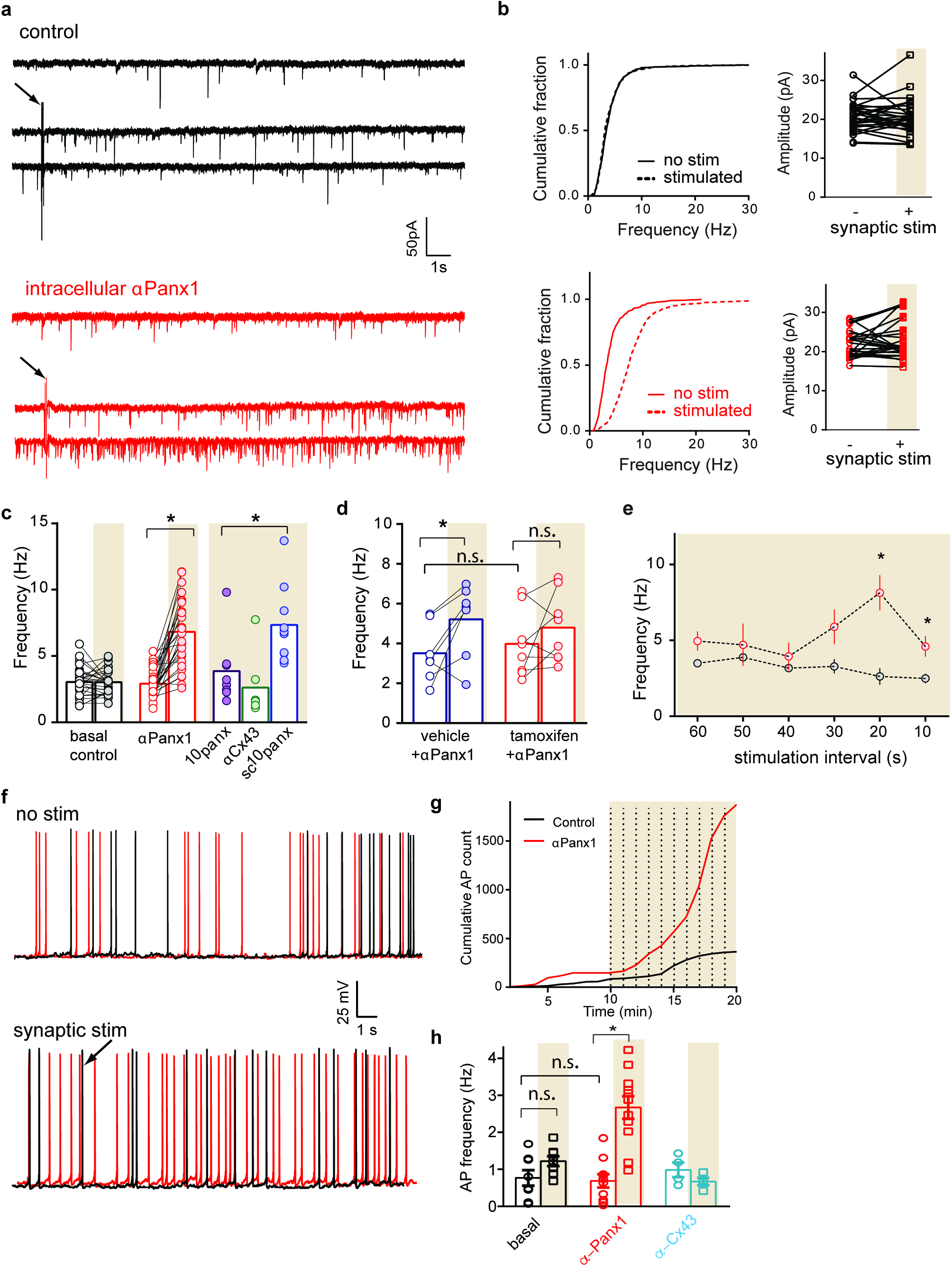
Block of postsynaptic pannexin-1 augments stimulation induced glutamate release and action potential frequency. **a)** Exemplar sweeps from CA1 pyramidal neurons, voltage clamped at -70 mV under control conditions (top) and after paired pulse stimulation of Schaffer collaterals (arrow). The red traces are from a neuron with 0.25ng/μl αPanx1 (Panx1 blocking antibody) in the patch pipette, without (top red trace) and with synaptic stimulation (arrow). **b)** cumulative probability distributions of EPSP frequency (black lines are control and red lines are with postsynaptic αPanx1. The plots on the left show the paired distribution of EPSP amplitudes, which were not significantly different (Wilcoxon matched-pairs signed rank test, control (black) p=0.83, with αPanx1 (red) p=0.3). **c)** Histogram showing the frequency of excitatory postsynaptic events before and after synaptic stimulation (shaded regions). Note the significant (Kruskal-Wallis test with Dunn’s post hoc) increase in the frequency of events following stimulation, indicating prolonged release. Two blockers of Panx1, αPanx1 (in the pipette, p<0.0001) and ^10^panx (100 μM in the bath, p<0.0001) caused prolonged release following stim. The control for αPanx1 (αCx43, p=0.084) and for ^10^panx (sc^10^panx, p=0.074) were not effective. **d)** The prolonged glutamate release induced by αPanx1 was present in Panx1^fl/fl^ mice treated with vehicle (left bars) but not in knockout mice (treated with tamoxifen; right bars). Kruskal-Wallis with Dunn’s post hoc: vehicle control vs. vehicle stim, p<0.05; vehicle control vs. tamoxifen control, p>0.05; tamoxifen control vs. tamoxifen stim, p>0.05). **e)** Delivery of paired pulse synaptic stimulations at the indicated time intervals reveals a stimulation-response curve for asynchronous release when αPanx1 is in the pipette (red circles). **f)** Overlaid one-minute current clamp (V_m_ = – 54mV) recordings, showing spontaneous action potentials in control (black) and neurons with αPanx1 in the pipette (red). Upper recordings did not receive synaptic stimulation. Lower recordings received paired pulse (20s intervals) at the arrow. Note that more action potentials are generated with αPanx1 is in the pipette and Schaffer collaterals are stimulated. **g)** plot of the cumulative number of action potentials over a 20 minute period. Synaptic stimulation (vertical dashed lines) dramatically increases total number of action potentials when αPanx1 is in the pipette. **h)** Comparison of the mean action potential frequency with the Kruskal-Wallis test with Dunn’s post hoc on mean number of action potentials calculated for 5 minutes at the beginning and end of the recordings. Control vs. control with stim, p=0.035; control vs. αPanx1 baselines, p=0.7; control with stim vs αPanx1 with stim, p=0.0048. *p = <0.05 data is represented as mean±SEM

Responses to postsynaptic delivery of αPanx1 were evaluated in our pyramidal neuron specific, conditional Panx1 knockout mice (Panx1^-/-^) ^18^. The control animals received vehicle alone (5% ethanol/95%corn oil) and showed Schaffer collateral stimulation-dependent prolonged synaptic events when αPanx1 was in the pipette (Fig. 1c). Prolonged release did not occur in Panx1^-/-^ neurons (tamoxifen treated Panx1^fl/fl^-*wfs1*-Cre mice) when αPanx1 was in the pipette (Fig 1c). The lack of an effect of αPanx1 in Panx1^-/-^ mice demonstrates specificity of the antibody ^17^ and specificity of the stimulation-induced prolonged release for postsynaptic Panx1.

We further examined the specificity of this novel synaptic activity on Panx1 with a peptidergic blocker, ^10^panx (100 μM) and its scrambled peptide control (sc^10^panx; 100 μM) ^24^. Bath applied ^10^panx paired with afferent stimulations resulted in a similar prolonged synaptic activity to αPanx1 (n=6 neurons, duration = 9.9±1.5 s), but sc^10^panx did not (Fig. 1b). Thus, three distinct ways of eliminating Panx1 activity in either single postsynaptic CA1 neurons (αPanx1) or *en masse* in the slice (^10^panx or Panx1^-/-^) resulted in prolonged glutamate release when Schaffer collaterals were stimulated.

Different inter-stimulation intervals were delivered to generate a stimulation-response curve (Fig. 1d). Significant (paired t-test; p<0.05) augmented release was observed with intervals of 10 and 20 s (0.17 and 0.33 Hz, respectively), but not at longer or shorter intervals (Fig. 1d). While 20s between afferent stimulations appeared optimal, both single (one pulse) and minimal (one pulse with a 50% failure rate) stimulations were also effective (Fig. S2a). Finally, the aPanx1 had to be delivered intracellularly to be effective (Fig. S2b).

### Block of postsynaptic Panx1 increases action potential frequency following stimulation

We predicted that this prolonged glutamate release would increase neuronal excitability ^29^. To test this, we recorded action potentials in CA1 neurons in response to afferent stimulation when αPanx1 was in the pipette. Figure 1f shows 10 s recordings of action potentials from CA1 neurons in current clamp, with E_m_ held at -54 mV, which is below the average threshold (-47 mV) for these cells ^30^. Without Schaffer collateral stimulation, inclusion of αPanx1 in the pipette did not significantly change action potential frequency (n=10, paired t-test, p=0.77). When postsynaptic αPanx1 was combined with paired-pulse stimulation, as in the synaptic recordings presented above, increased action potential frequency occurred (Fig. 1fg). There was a substantial increase in the cumulative number of action potentials generated over a 10-minute stimulation period (Fig. 1f). Thus, the data in Figure 1 show that block of postsynaptic Panx1 augments excitability following afferent stimulation via asynchronous glutamate release.

### TRPV1 is required for prolonged release and augmented excitability

The stimulation induced asynchronous release during Panx1 block reminded us of transient receptor potential vanilloid 1 (TRPV1) channels in the NTS, where their activation by anandamide (AEA) causes asynchronous glutamate release ^13,31^, even though our effect was substantially longer. We hypothesized that TRPV1 channels could account for the prolonged release observed when Panx1 is blocked. The expression of TRPV1 in the hippocampus is controversial and several groups either support ^32,33^ or refute ^34^ its presence. Therefore, we used immunoelectron microscopy in wildtype and TRPV1 knockout (TRPV1^-/-^) mice to investigate if TRPV1 is expressed at synapses in the CA1. We found that ~20% of terminals were TRPV1 positive, and immunogold labelling was almost undetectable in the TRPV1^-/-^ mice (Fig 2ab).

**Fig. 2.**
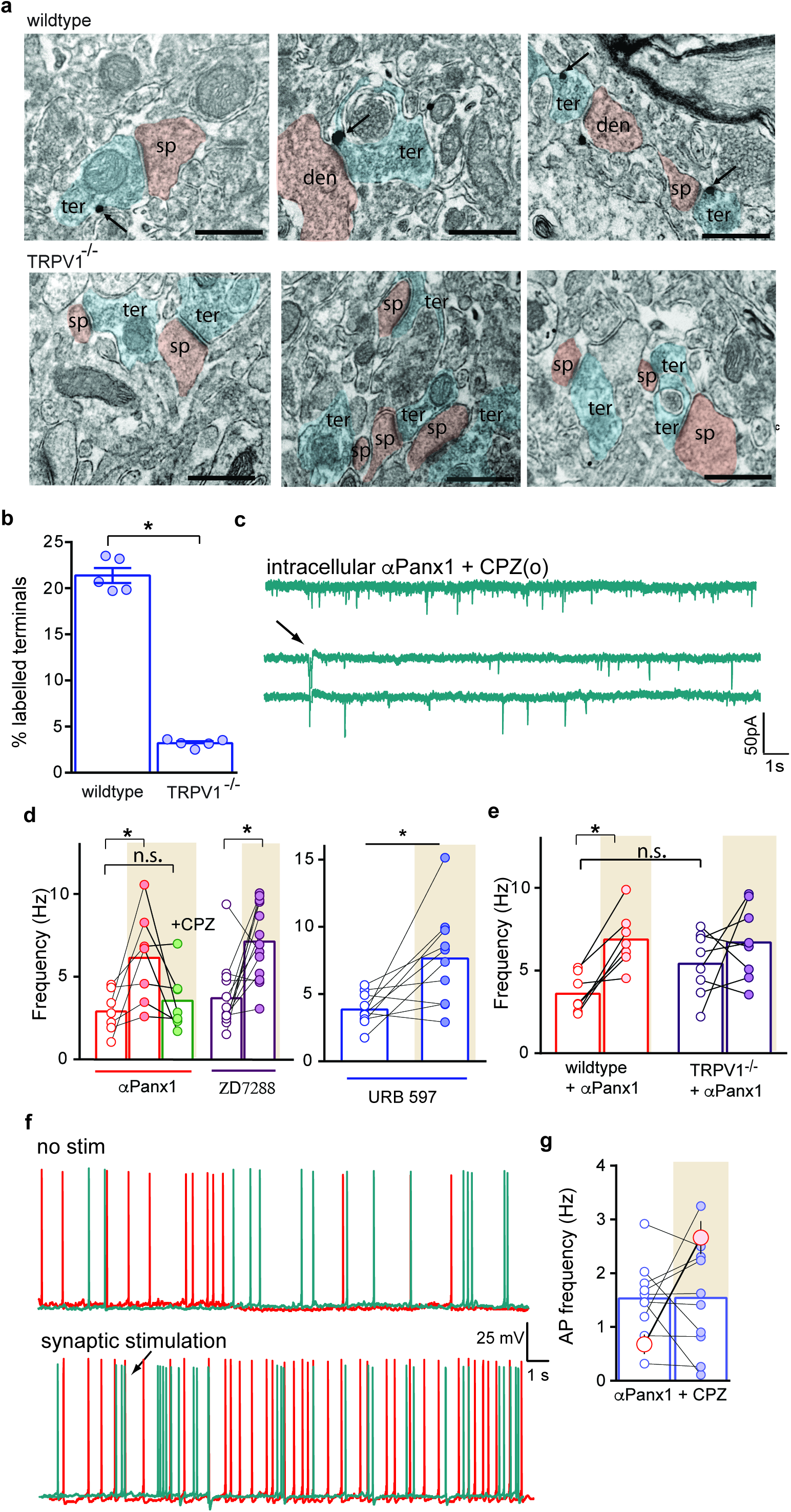
TRPV1 is required for postsynaptic Panx1 block-induced prolonged neurotransmission and excitability. **a)** Immunogold labelling of TRPV1 labelling in wildtype (top) and TRPV1^-/-^ knockout (bottom) mice in area CA1. Arrows indicate TRPV1 immunogold particles. Terminals are blue and CA1 spines or dendrites red. Scale bars represent 1 μm. **b)** Comparison of the percentage of labelled terminals (each point is an individual animal) showed a significantly fewer number of TRPV1 positive terminals in the TRPV1^-/-^ mice (Mann Whitney test, p=0.079). **c)** The αPanx1 and synaptic stimulation (arrow, paired pulse at 20s intervals) prolonged release was blocked by bath applied TRPV1 antagonist, capsazepine (CPZ; 10 μM). **d)** Comparison of the sEPSC frequency with TRPV1 blocked (CPZ) or Ih blocked (ZD7288 (10 μM)). Kruskal-Wallis with Dunn’s post hoc: αPanx1 baseline vs. αPanx1 with stim, p=0.014; αPanx1 baseline vs. αPanx1 + CPZ with stim, p>0.9; ZD7288 baseline (open purple circles) vs. ZD7288 with stim (closed purple circles), p=0.01. Comparison of the FAAH inhibitor, URB597 showed a significant (p=0.01; Wilcoxon paired) increase upon stimulation. **e)** Comparison of the changes in sEPSC frequency in wildtype (C57BL6/J) compared to TRPV1^-/-^ mice. Kruskal-Wallis with Dunn’s post hoc: wildtype baseline vs. wildtype with stim, p=0.024; wildtype baseline vs. TRPV1^-/-^ baseline, p=0.53; TRPV1^-/-^ baseline vs. TRPV1^-/-^ with stim, p>0.9. **f)** Current clamp recordings of action potentials with αPanx1 in the pipette (red) and CPZ in the bath (green). Synaptic stimulation (lower traces, arrow) increased action potential frequency and this was blocked by CPZ. **g)** Comparison of the action potential frequency with (shaded) and without synaptic stimulation in the presence of CPZ (bars; blue points) showed no significant difference (Wilcoxon matched pairs, p=0.84). The red circles show the change in frequency when CPZ was not in the pipette (data from Fig. 1e). *p = <0.05 data is represented as mean±SEM

If activation of TRPV1 is responsible for prolonged glutamate release during intracellular delivery of αPanx1 to postsynaptic CA1 neuron then blocking or knockout of TRPV1 should abolish prolonged release. Addition of 10 μM capsazepine (CPZ) to the bath, while blocking postsynaptic Panx1, prevented afferent stimulation-induced asynchronous release (Fig 2cd) and increased action potential frequency (Fig. 2ef). CPZ is reported to alter activity of Ih in pyramidal neurons ^35^ so we evaluated the specific Ih blocker, ZD7288 (10 μM), which failed to prevent prolonged release (Fig 2d). CPZ did not affect the basal rate of spontaneous events in TRPV1^-/-^ mice compared to wild type animals (wildtype frequency was 2.9±0.5 Hz versus 3.9±0.7 Hz in TRPV1^-/-^ slices; p=0.45, Mann Whitney U test, n=7 and 7, respectively).

We inhibited fatty acid amid hydrolase (FAAH) by addition of 1 μM URB597 to the bath, which increases AEA concentration and reproduced the stimulation-dependent increase in neurotransmission (Fig. 2d), suggesting a key role for AEA. TRPV1^-/-^ mice did not show an effect of postsynaptic αPanx1 during afferent stimulation, but wild type mice had robust stimulation-induced prolonged release (Fig. 2g). Finally, bath application of capsaicin (CAP; 1 μM) to slices did not alter spontaneous release when compared to controls, but interestingly, with concomitant Schaffer collateral stimulation in the presence of CAP there was increased EPSP frequency. (Control mean frequency = 3.6±0.4 Hz (n=7), with CAP = 3.7±0.6 Hz (n=3), which were not significant (p>0.05) from each other. Frequency with CAP+stimulation = 7.1±0.6 Hz (n=13), which was significantly increased versus CAP without stimulation at p>0.05; Kruskal-Wallis test, p=0.0.0009, H=18.81). This suggests that stimulation may induce insertion of TRPV1 into the plasma membrane, which may be regulated by phosphorylation of the channel ^36^. Together, these data suggest that TRPV1 in the hippocampus may be responsible for prolonging glutamate release when Panx1 is blocked.

### Panx1 regulates tissue AEA levels

How is TRPV1 activated when postsynaptic Panx1 is blocked? Since AEA is an endogenous ligand of TRPV1, and AEA has an important role in asynchronous release in the NTS ^13^, we reasoned that Panx1 could be regulating its synaptic concentration. Tissue concentrations of AEA were quantified using mass spectrometry (Fig. 3a). Basal levels were 7.3±0.8 pg/mg (n=14 hippocampal slices from 6 rats). This significantly increased to 11.9±1.2 pg/mg when Panx1 was blocked with ^10^panx (p=0.002 vs control; one-way ANOVA, n=13 slices from 6 rats). It was reported previously that TRPV1 may regulate AEA transport ^37^. However, CPZ (10 μM) did not change tissue AEA levels (with CPZ, AEA = 7.4±0.7 pg/mg; n=8 slices from 4 rats; p>0.05, one-way ANOVA). These data support the idea that Panx1 block increases total AEA concentration in hippocampal slices.

**Fig. 3.**
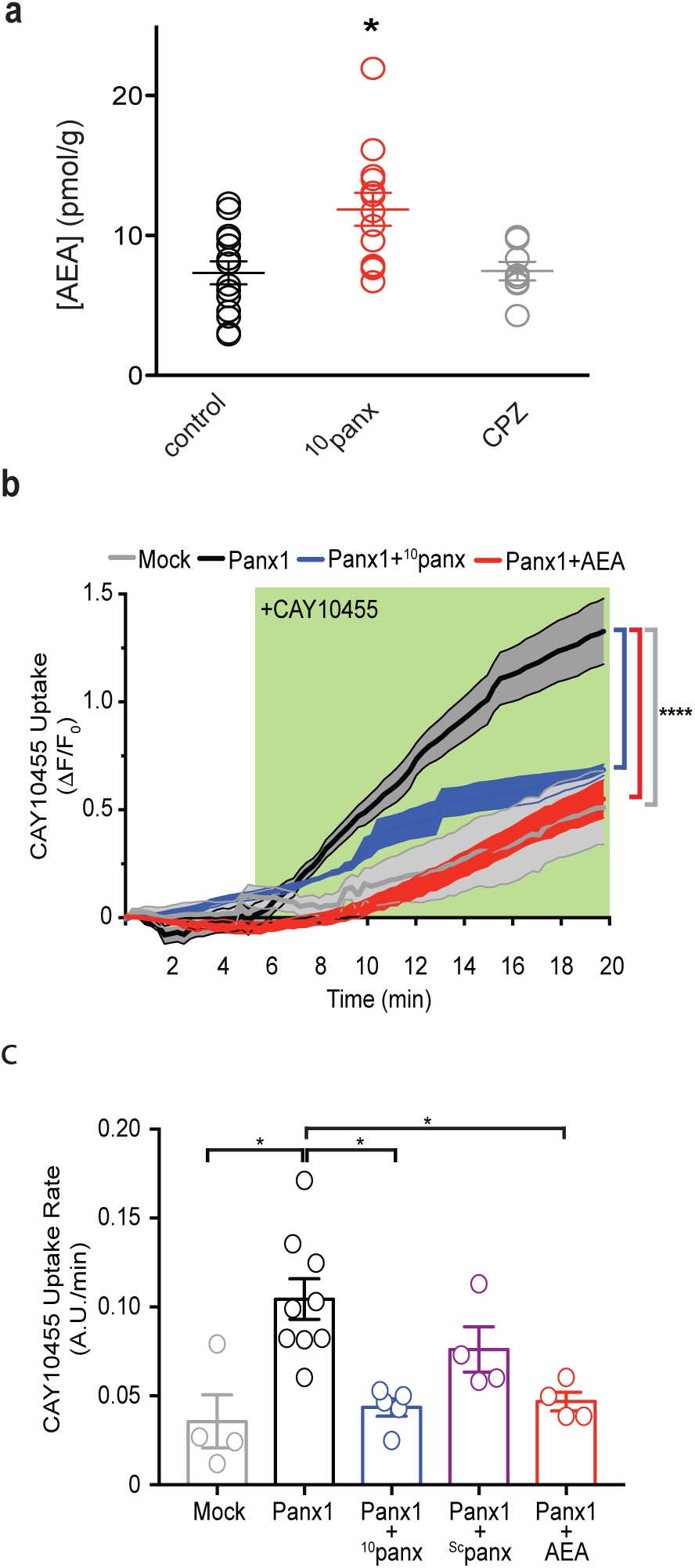
Block of Panx1 increases tissue levels of anandamide and ectopic expression promotes uptake of a fluorescent anandamide derivative. **a)** The Panx1 blocker, ^10^panx was bath applied to acute hippocampal slices and AEA content quantified by mass spectrometry. Panx1 block, but not TRPV1 block (with CPZ) increases the concentration of AEA. Kruskal-Wallis with Dunn’s post hoc: control vs. ^10^panx, p<0.05; control vs. CPZ, p>0.05. **b)** Panx1 expression in HEK293 cells augmented uptake of fluorescent anandamide (5μM CAY10455) compared to mock transfected controls. The green box indicates the presence of CAY10455 in the bath. CAY10455 uptake was inhibited by the Panx1 antagonist, 100μM ^10^panx or unlabelled AEA (50μM). Two-way ANOVA revealed significantly (p<0.05) different increases over time as indicated. **c)** Comparison of the CAY10455 uptake as an average of the fluorescence between 15–20 min. The scrambled control for ^10^panx, sc^10^panx did not prevent CAY10455 uptake. One-way Anova with Tukey’s multiple comparisons: mock transfected vs. Panx1 transfected p=0.005; Panx1 vs ^10^panx p=0.019; Panx1 vs AEA p=0.007; all other comparisons were not significant. *p = <0.05 data is represented as mean±SEM.

### Ectopic expression of Panx1 in HEK293T cells augments fluorescent AEA uptake

One possible mechanism for Panx1 block to increase tissue AEA concentration is by Panx1 facilitating clearance (uptake) of AEA. If true, ectopic expression of Panx1 should augment the uptake of the AEA analogue, CAY10455, which fluoresces only after esterase cleavage in the cytosol. This assay was chosen over the [^3^H]AEA transport assay because radiolabelled AEA measurements do not distinguish between uptake and membrane accumulation. Transient transfection of Panx1 in HEK293T cells increased uptake of bath applied CAY10455 compared to mock transfected controls (5 μM; Fig 3b). This uptake was blocked by ^10^panx, but not sc^10^panx (Fig 3c). Importantly, a 10-fold excess of unmodified AEA (50 μM) prevented CAY10455 influx into Panx1 expressing HEK293T cells (Fig 3c). While 50 μM is a high concentration of AEA it was required to quantify block of CAY10455 fluorescence because lower concentrations of CAY10455 were dim. Thus, our data presented in Fig 3 support a model whereby extracellular AEA clearance is facilitated by Panx1 channels.

### AEA blocks dye flux through Panx1

How could Panx1 facilitate AEA uptake? One possibility is that these non-selective ion / metabolite channels are permeable to AEA and therefore are functioning as a synaptic AEA transporter. AEA is an uncharged molecule, making it impossible to measure flux (as current) with electrophysiology. However, Panx1 channels flux molecules < 1kD, so we reasoned if the 0.35 kD AEA is permeable it should compete with dye flux through the channel and this would be a sensitive assay for AEA influx. We modified the cell-attached patch clamp to include the Panx1 permeable dye, sulforhodamine 101 (SR101; 0.6kD) and the Panx1 impermeable dye, FITC-dextran (3-5kD) in the pipette ^24^. Voltage-dependent activation of Panx1 ^38^ in cultured hippocampal neurons induced single channel currents and SR101 uptake (Figs 4acd). FITC-dextran uptake occurred only in the whole-cell configuration (Fig 4b), indicating its occlusion is a control for membrane integrity in the cell-attached mode. The Panx1 blocker, ^10^panx (100μM) in the pipette prevented SR101 influx and ionic currents (Fig. 4cd). Similarly, 50μM AEA in the pipette reduced SR101 influx (Fig 4cd).

**Fig. 4.**
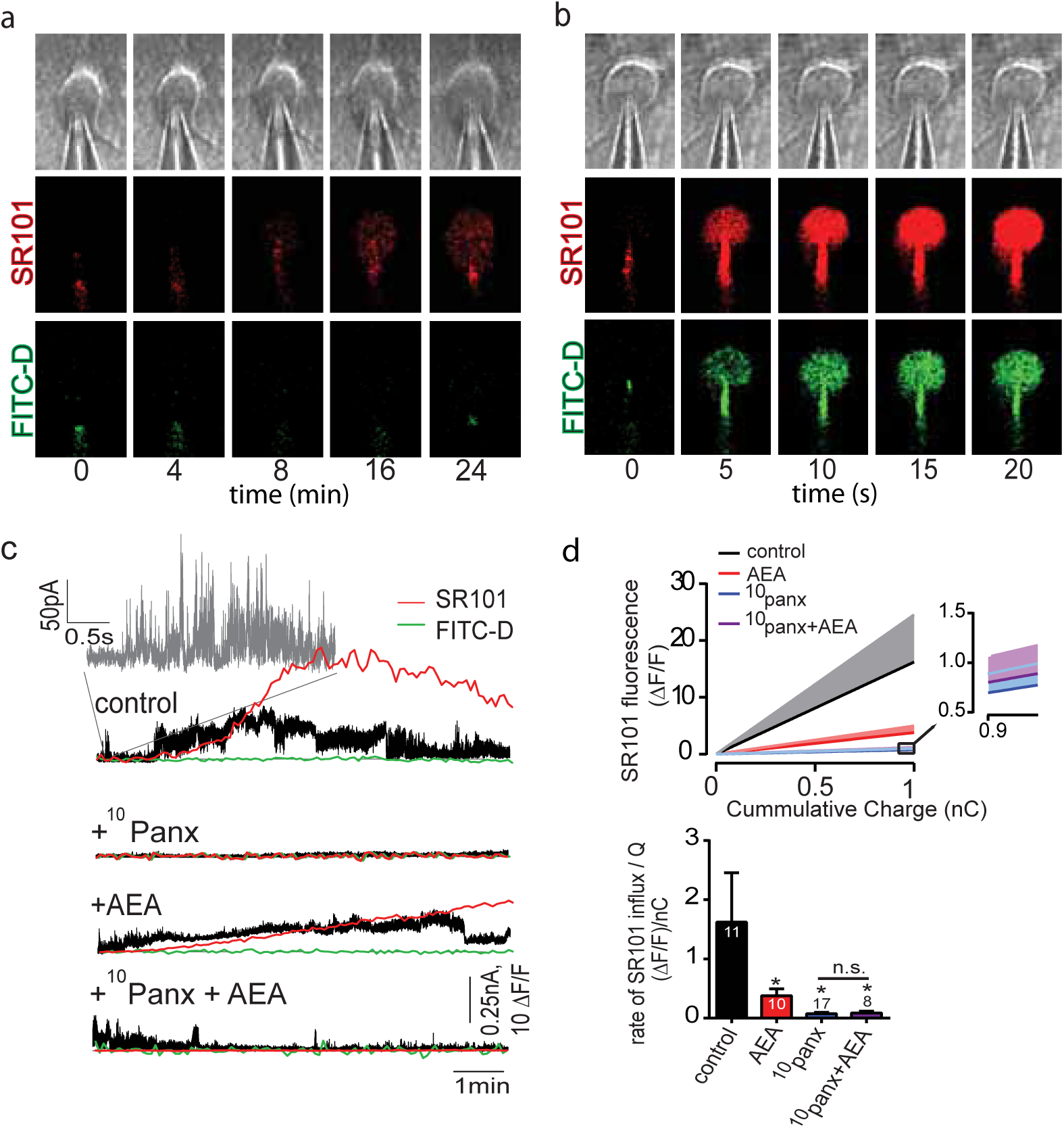
Anandamide competes with dye uptake through pannexin-1 in neurons. **a)** Cultured hippocampal neurons were patch-clamped in the cell-attached mode with SR101 (red) and FITC-dextran 4kDa (green) in the pipette in addition to a cocktail of blockers for other channels / receptors. Only Panx1 permeable SR101 enters neurons during voltage ramp depolarizations. **b)** Break-in to whole-cell mode leads to rapid loading of all dyes. **c)** Superimposed current recordings and dye flux. The inset (grey trace) shows the typical fast gating of pannexin-1 single channels. SR101 influx occurred when channel activity was detectable in the patch. Note that the pannexin-1 impermeable FITC-dextran (green) did not enter neurons. With 100μM ^10^panx (2^nd^ trace) or 50μM AEA (3^rd^ trace) in the pipette, SR101 influx was attenuated. The combined addition of AEA and ^10^panx (bottom trace) also reduced SR101. **d)** Comparison of the rate of change in SR101 fluorescence in neurons (top) and the rate of SR101 influx normalized to total charge (Q) (bottom). Kruskal-Wallis test with Dunn’s multiple comparisons: control vs AEA, p=0.035; control vs ^10^panx p<0.0001; control vs ^10^panx+AEA, p=0.006; all other comparisons p>0.05.

### Constitutively active Panx1 facilitates AEA flux

Panx1 may act to facilitate AEA flux by being constitutively active or through recruitment during synaptic activity or both. If there is basal Panx1 activity, we predicted that AEA loaded into CA1 neurons in slices via the patch-pipette could efflux the cell and activate presynaptic TRPV1 without stimulation. This would therefore show that AEA can have bidirectional flux and is consistent with models of AEA transport ^39^. CA1 pyramidal neurons were loaded with 50 μM AEA via the patch pipette and spontaneous excitatory synaptic currents (sEPSC) were recorded. Postsynaptic AEA increased sEPSC frequency without the requirement for afferent stimulation (Figs. 5abef). The increase in sEPSC frequency occurred 3.6±1.1 min after whole-cell formation in 8 of 10 cells tested, with a range of 19 s – 9 min. As shown in Fig. 5, the stimulation independent increase in sEPSC frequency with postsynaptic loading of AEA was blocked by either αPanx1 in the pipette (Fig. 5c) or CPZ in the bath (Fig 5d). When Panx1 was blocked with αPanx1 in the presence of postsynaptic AEA, afferent stimulation induced prolonged synaptic events (Fig. 5f) similar to those in Fig. 1, indicating that afferent stimulation induced AEA was not likely normally coming from postsynaptic CA1 neurons via Panx1 despite the ability of Panx1 to efflux AEA when neurons were loaded with AEA.

**Fig. 5.**
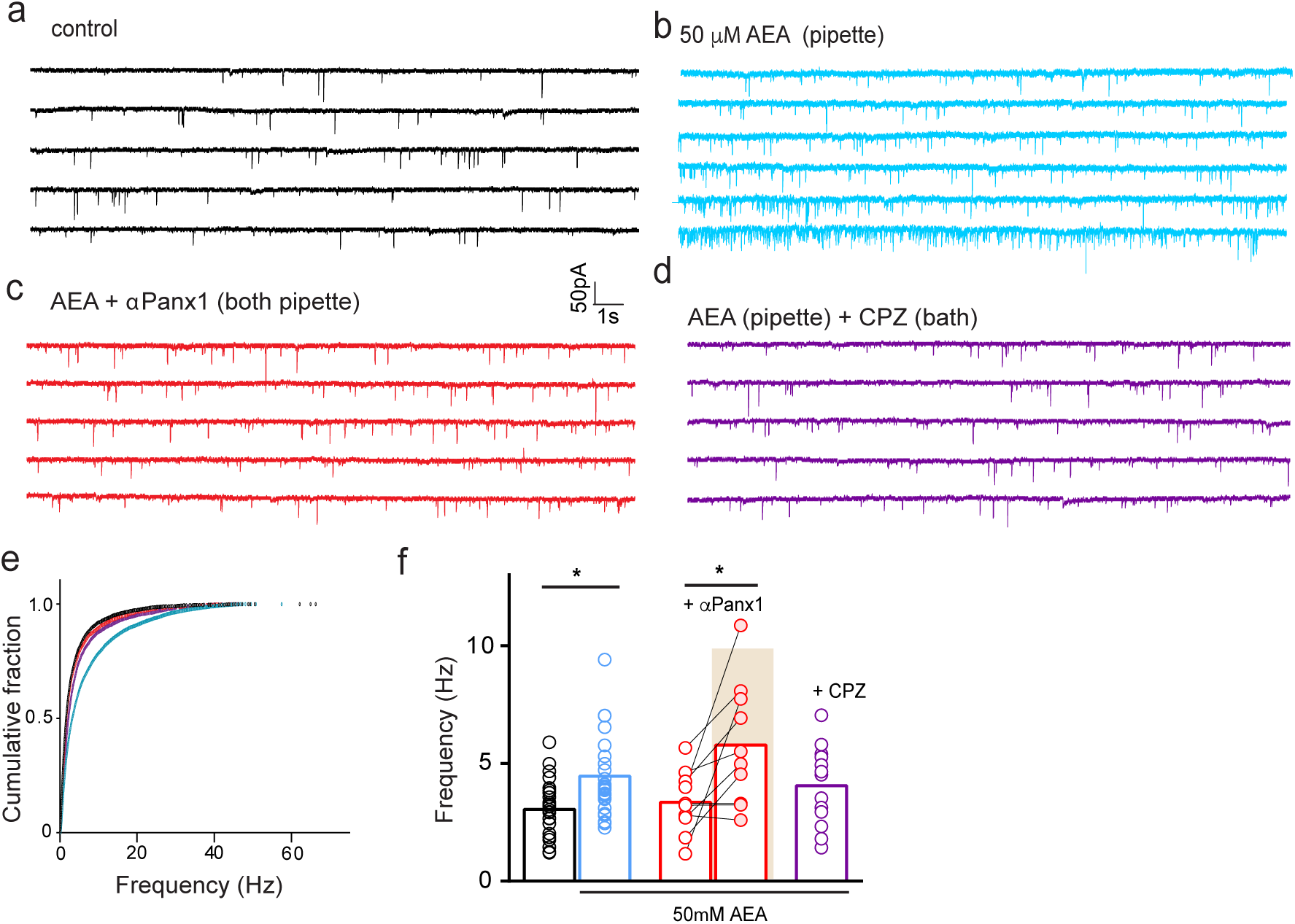
Loading of AEA in the postsynaptic neurons augments glutamate release in a Panx1 and TRPV1 dependent manner. **a-d)** Recordings from 4 different CA1 neurons under the conditions indicated. Note that AEA in the pipette (b) increases sEPSC frequency with a delay after break-in. Either αPanx1 in the pipette (c) or CPZ in the bath (d) block this increase. **e)** Cumulative probability plot of the instantaneous frequency of sEPSCs. Colors of the lines match the same conditions shown in f. **f)** Comparison of the mean±SEM frequency of sEPSC events. Kurskal-Wallis test with Dunn’s post hoc: control (black) vs. AEA (blue), p=0.017; control vs. αPanx1 baseline (red open circles), p>0.9; αPanx1 baseline vs αPanx1 with stimulation, p=0.006; control (black) vs AEA+CPZ (purple), p=0.515. *denotes significantly different.

### Asynchronous release does not involve postsynaptic Ca^2+^

The requirement for afferent stimulation during application of postsynaptic αPanx1 suggests that AEA production may occur in a Ca^2+^-dependent way ^40^, as is the case for most known retrograde signals ^41^. These typically regulate synchronous neurotransmitter release by altering the coupling of voltage-gated Ca^2+^ channels to release machinery ^41^. This is unlikely in the present study because with postsynaptic αPanx1 the paired-pulse ratio (PPR) was unchanged (Fig. S3). We argue, based on Fig 5, against a postsynaptic source of AEA because prolonged release occurred when AEA was loaded into the postsynaptic neuron and Panx1 was blocked. However, if AEA is being produced in the CA1 neuron, it would likely require increased postsynaptic Ca^2+^ for production ^40^. When 10 mM BAPTA and αPanx1 were added to the pipette to chelate postsynaptic Ca^2+^, prolonged glutamate release upon Schaffer collateral stimulation was still evident (Fig. 6ac). In contrast, incubation of hippocampal slices in membrane permeable EGTA-AM to chelate Ca^2+^ in all cells (50μM; minimum 15 min loading time) blocked stimulation-induced prolonged release (Fig 6ac). We chose EGTA-AM over BAPTA-AM for bulk loading because EGTA does not block synchronous release, but is sufficient to inhibit the slower increases in Ca^2+^ required for asynchronous release ^42^. AEA can activate cannabinoid receptors (i.e. CB1). Therefore, we used the CB1 receptor inverse agonist, AM251 (3 μM) and ruled out roles for these pathways in Panx1 modulated prolonged release (Fig. 6bd). As a positive control for AM251, we isolated inhibitory postsynaptic GABA currents (IPSC) by removing picrotoxin and addition DNQX to block AMPA receptors while adding αPanx1 to the CA1 neuron via the patch pipette. As shown in Fig S4, there was a progressive inhibition of IPSCs that was reversed by AM251, indicating that blocking Panx1 can activate CB1 receptors at GABA synapses onto CA1 neurons and suppress inhibition.

**Fig. 6.**
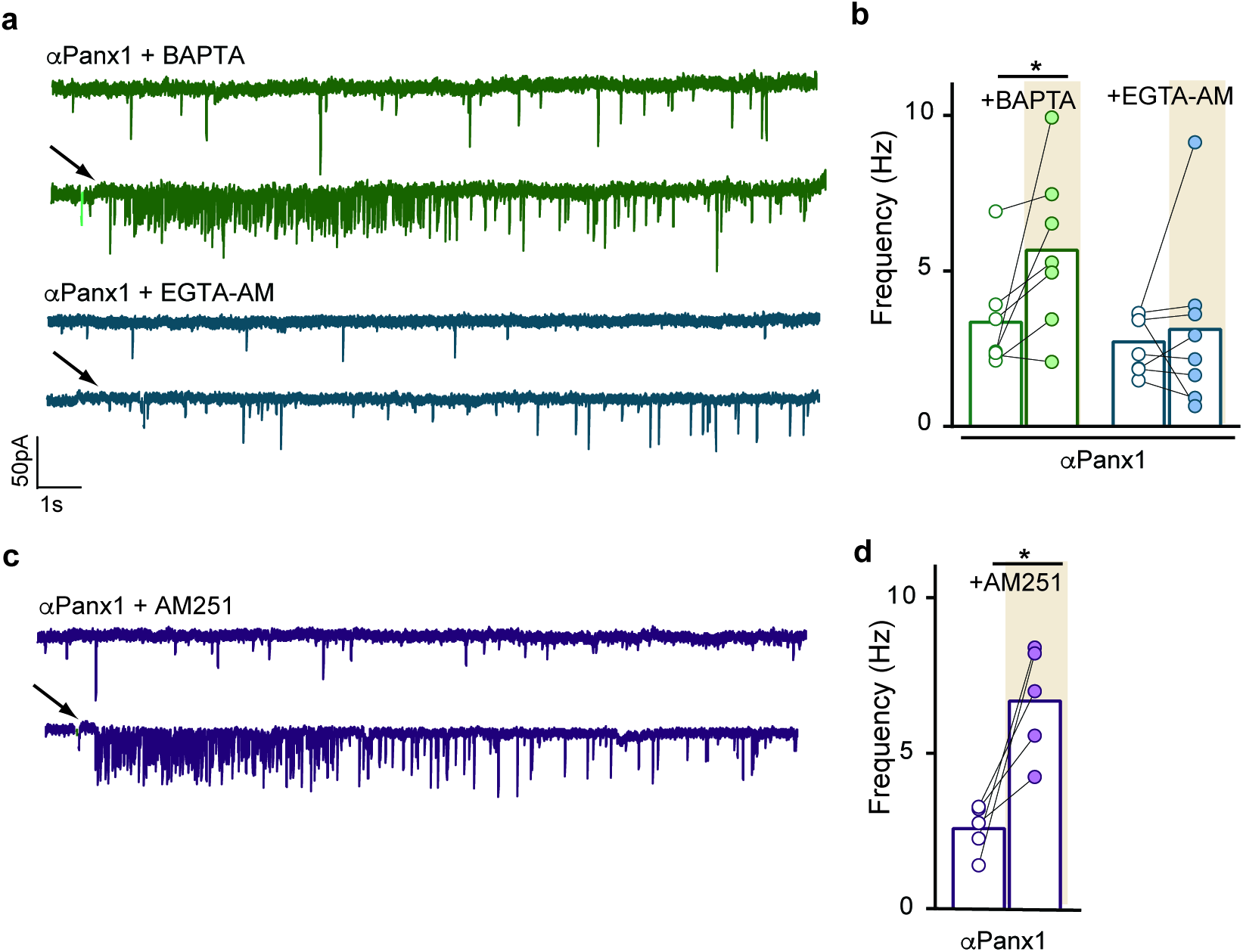
Prolonged neurotransmission does not require postsynaptic Ca^2+^ or CB1 receptors. All recordings contained αPanx1 in the pipette. **a)** Exemplar recordings from 2 different CA1 neurons under high postsynaptic Ca^2+^ buffering conditions (top; BAPTA) and low Ca^2+^ buffering conditions in all cells (blue; EGTA-AM). The slow Ca^2+^ buffer prevented synaptic bursting in response to afferent stimulation (arrows), but strong buffering in the postsynaptic cell did not. **b)** Comparison of the frequency changes in glutamate neurotransmission with different Ca^2+^ buffers. Wilcoxon matched pairs test: BAPTA baseline vs. BAPTA stimulated, p=0.031; EGTA control vs. EGTA stimulated, p=0.5. **c&d)** The classical retrograde transmitter receptor, CB1 R was ruled out by application of its specific antagonist, AM251. Wilcoxon matched pairs test p=0.03. Bars show mean. *Denotes significance at p>0.05.

### Panx1 knockout augments electrical kindling in a TRPV1 dependent manner

The prolonged glutamate release and suppression of GABA synapses described above should contribute to enhanced excitability *in vivo*. We investigated a potential role for Panx1 and TRPV1 in epileptogenesis using the electrical kindling model (Fig 7a). Electrical kindling stimulation of the dorsal hippocampus via chronically implanted bipolar electrodes induced seizures, which we quantified with electrographic recordings (i.e. Fig 7b) and by assignment of seizure severity on the Racine scale ^43^ for each session. Panx1^-/-^ mice required fewer kindling sessions to become fully kindled (i.e. 3 stage 5 seizures) (Fig. 7c) compared to untreated wild type mice and wild type mice that received tamoxifen (WT n=5; WT^Tam^ n=5; Panx1^-/-Tam^ n=6; one-way ANOVA [WT^Tam^ vs Panx1^-/-Tam^ p=0.0091]). Consistent with fewer sessions to reach 3 stage 5 seizures, Panx1^-/-^ mice kindled more quickly than both control groups (Fig. 7d).

**Fig. 7.**
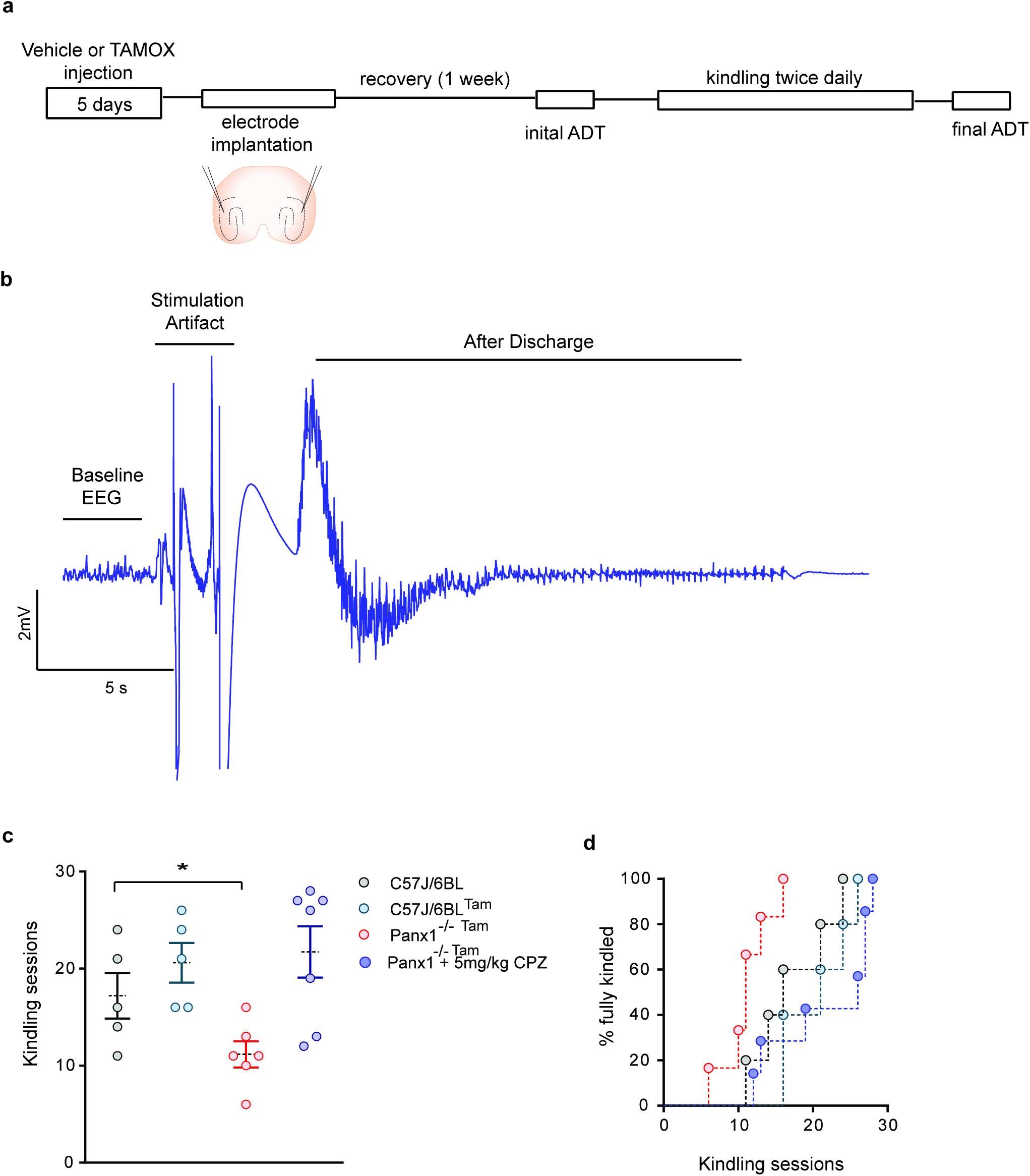
Panx1 knockout augments epileptogenesis in a TRPV1-dependent manner. **a)** Electrical kindling paradigm. Mice received electrical kindling stimulation in the ventral hippocampus twice daily until fully kindled, indicated by 3 consecutive stage 5 seizures**. b)** Example EEG recording showing the ictal event (afterdischarge). **c)** The number of kindling sessions required to reach 3 stage 5 seizures was significantly lower in Panx1^-/-^ mice and this was reversed by i.p. injection of CPZ prior to each kindling session. One-way ANOVA with Tukey’s post hoc: control (black) vs Panx1^-/-^ p=0.012. Control vs. Panx1^-/-^ + CPZ (purple) p>0.05. **d)** Comparison of the time course of kindling reveals that Panx1^-/-^ mice reached stage 5 seizures sooner than the controls. This was reversed by CPZ.

If this enhanced epileptogenesis in Panx1^-/-^ was due to increased TRPV1 activation, as the *in vitro* data indicate, then intraperitoneal (i.p.) injection of CPZ prior to each kindling session should return the kindling rate to the control level. 30 minutes prior to each kindling session, Panx1^-/-^ mice received an i.p. injection of 5 mg/kg CPZ. In the presence of CPZ, the kindling rate was not different from the control (Fig. 7cd; CPZ n=7; one-way ANOVA [control vs. CPZ p=0.4878]; [Panx1^-/-Tam^ vs. CPZ p=0.0104).

## Discussion

Here we report a novel form of prolonged synaptic glutamate release onto CA1 neurons that is dependent upon postsynaptic Panx1, presynaptic TRPV1, AEA and afferent stimulation. Several Panx1 blockers, including a validated blocking antibody directly delivered to single postsynaptic neurons, and conditional genetic deletion of Panx1 caused prolonged (seconds) glutamate release in response to Schaffer collateral stimulation. There was a parallel robust increase in action potential generation. We propose that postsynaptic Panx1 channels facilitate rapid clearance of afferent stimulation-produced AEA so that when the channels are closed, AEA accumulates and acts at TRPV1 to induce glutamate release (see Fig. S5). As a consequence of increased glutamate there was enhanced network synchrony seen as faster epileptogenesis during kindling. Importantly, augmented epileptogenesis was reversed by *in vivo* administration the TRPV1 antagonist, CPZ.

### TRPV1 and prolonged glutamate release

In the hippocampus, short-term alterations of synaptic function are most often seen at inhibitory synapses and manifest as prolonged periods of transmitter release following stimulation ^44^. An interesting example is the activation of P2X2 receptors at CA1-interneuron synapses, leading to asynchronous glutamate release ^12^. While it has been controversial whether TRPV1 is broadly expressed in the hippocampus, it has been described in specialized Cajal-Retzuis cells ^34^. Furthermore, there are several functional demonstrations of TRPV1 activity in the brain. In hippocampal mossy fibres TRPV1 can induce a postsynaptic Ca^2+^-dependent LTD of both GABAergic and glutamatergic signalling via receptor internalization^45,33^. In addition, we show here that Panx1 block can activate CB1 receptors on GABAergic neurons to suppress IPSCs, which would be expected to contribute to the overall excitation of the system. It is not clear if this was AEA-dependent and future studies will explore this possibility.

We have discovered that TRPV1 can induce prolonged glutamate release through a postsynaptic Ca^2+^-independent mechanism because high concentrations of BAPTA in the pipette failed to prevent prolonged release. It is possible that AEA spillover from adjacent neurons lead to prolonged glutamate release during postsynaptic BAPTA loading. We do not however favour this possibility because open Panx1 channels in these adjacent cells likely rapidly clear AEA. The simplest explanation for our data is that TRPV1 is expressed in presynaptic compartments that synapse with CA1 neurons and that TRPV1 is activated by Ca^2+^-dependent production of the endovanilloid, AEA. TRPV1 could be present in CA3 axons^46^, or intriguingly, in Cajal-Retzius cells ^34^. We report TRPV1 immunogold labelling in ~20% of presynaptic compartments in the CA1 region, which was absent in TRPV1^-/-^ mice. However, we cannot determine if TRPV1 is in CA3 terminals or novel synapses from Cajal-Retzius cells. It is important to note that other, unidentified compartments were also labelled for TRPV1 in our immunohistochemical electron microscopy; the function of TRPV1 in these areas in not known.

### Panx1 regulates AEA concentration and TRPV1 activation

AEA is a well-characterized ligand for TRPV1 ^47^ with reported EC50 values between 0.7 – 5 μM in expression systems and ∼10 µM in DRG neurons ^48^. Thus, AEA would need to reach µM levels in synapses to effectively activate TRPV1. We show here that blocking Panx1 increased the concentration of AEA in hippocampal slices and propose that this increase in AEA is sufficient to activate TRPV1. Although reported bulk levels of AEA in brain are low (44 pmol/g)^49^, the requirement for Schaffer collateral stimulations to induce TRPV1-mediated prolonged release is consistent with the concentration of AEA reaching transiently high levels. Excitatory neurons of the NTS release AEA to cause asynchronous glutamate release and enhanced excitation through recruitment of TRPV1 ^13^. In mossy fiber-CA3 synapses and DRG neurons the mechanism of TRPV1 activation by AEA is proposed to be intracellular (i.e. AEA is not released into extracellular space) ^45,50^, which is distinct from what we are reporting here.

What is the source of AEA leading to prolonged glutamate release onto CA1 neurons? The prevalent view is that AEA is a retrograde transmitter, synthesized in postsynaptic cells by Ca^2+^-dependent NAPE-PLD ^51^. Our evidence supports the notion that AEA is produced in a stimulation / Ca^2+^-dependent way to act at postsynaptic sites in the CA1 region. It is unlikely that this AEA is synthesized in the postsynaptic neuron and is acting as a retrograde transmitter because postsynaptic Ca^2+^ chelation failed to prevent asynchronous release when Panx1 was blocked. Additionally, we mimicked the stimulation evoked asynchronous release with bulk loading of AEA via the patch pipette, suggesting that Panx1 could facilitate release from the postsynaptic neuron. However, when this asynchronous release was prevented by blocking Panx1 it was evoked by Schaffer collateral stimulation. Together, this suggests that AEA biosynthesis is occurring in a different synaptic compartment than the CA1 neuron.

At excitatory CA3-CA1 synapses, the primary enzyme involved in AEA biosynthesis, NAPE-PLD, is reportedly expressed in presynaptic Schaffer collateral terminals ^52^. The AEA degrading enzyme, FAAH, in contrast is expressed in postsynaptic CA1 neurons ^53^. This molecular architecture is amenable to a stimulation and Ca^2+^-dependent mobilization of AEA from presynaptic terminals. It further implies that AEA should be transported into postsynaptic CA1 neurons for metabolism ^54^. Schaffer collateral stimulation-induced AEA release is unlikely to be facilitated by (putative) presynaptic Panx1 because bath applied blockers that act at extracellular sites on the channel mimicked the effect of intracellular application of postsynaptic blockers (i.e. increased asynchronous release). While the available data are most consistent with a presynaptic release of AEA, we cannot rule out other sources. There is emerging evidence that AEA can be synthesized and released from glial cells ^55,56^. So an alternative mechanism could be that Schaffer collateral stimulation drives mobilization of AEA from local glial cells, and Panx1 then facilitates clearance.

It is unconventional for AEA to signal in this proposed way – to be released presynaptically and cleared postsynaptically while its site of action is on the cell that is releasing it. We propose that Panx1, TRPV1 and AEA regulate glutamate release when a modulation of neuronal synchrony is required. For example, asynchronous release may promote both low and high frequency outputs from the hippocampus ^57^. Our identification of a role for Panx1 / TRPV1 during epileptogenesis (kindling) is likely a pathological manifestation of synchrony. It will be important in the future to determine if there are physiological roles for Panx1, AEA and TRPV1 in regulating hippocampal outputs during typical behaviour.

### Panx1 facilitates transport of AEA

Panx1 could facilitate removal of AEA in two ways: Firstly, by acting as the direct route of AEA flux across the membrane, or secondly, by regulating an unidentified transporter or AEA binding protein. The biophysical nature of AEA transport across membranes has been investigated and Panx1’s known properties are consistent with what we know about AEA transport ^58–60^: Because Panx1 is a channel, it would have a low temperature dependence for flux, and the Q_10_ for AEA transport is 1.4 ^59,60^. Panx1 is ubiquitously expressed and permeable to a broad range of ions and molecules. While it is not yet known how the weakly polar / lipophilic AEA could traverse the channel, Panx1 has hydrophobic amino acid residues in the pore ^61^ that could function as a scaffold for lipids to ‘jump’ independently of the water content. Alternatively, conformational changes or protonation / deprotonation of key pore lining amino acids could induce “dewetting” of the pore and a promote a hydrophobic environment. This ‘hydrophobic gating’ has been described in some bacterial ion channels ^62^. A further possibility is that AEA moves between Panx1 channels. Members of the gap junction superfamily, like Panx1, tend to cluster together and there could be hydrophobic routes for AEA between the channels, however this latter possibility is not consistent with the competition of AEA and fluorescent SR101 in single channels. It also remains possible that Panx1 can regulate the concentrations of other synaptically active molecules, such as ATP or additional endocannabinoids.

Alternative to flux through Panx1, the channels may directly or indirectly regulate the activity of an unidentified AEA membrane transporter. For example, it has been proposed that fatty acid binding proteins shuttle AEA (and lipids) within the cell to sites of degradation ^63^. Panx1 may localize closely with these proteins and facilitate binding of AEA. Another possibility is that Panx1 regulates FAAH activity, which has been reported to determine the concentration gradient for AEA influx ^64^ (see however ^65^). Regardless, we have shown here that Panx1 activity regulates the AEA concentration, leading to TRPV1 activation, prolonged glutamate release following afferent stimulation and augmented excitability that can contribute to epileptogenesis.

## Acknowledgments

The authors thank Dr. Keith Sharkey, Dr. Jaideep Bains, and Dr. Brian MacVicar for critical reading of the manuscript. The work was supported by grants from the Canadian Institutes of Health Research to MNH and RJT. Additional support was provided to RJT by the Cumming School of Medicine via the Ronald and Irene Ward Foundation and the Gwendolyn McLean Fund, and from the Hotchkiss Brain Institute. This work was supported by The Basque Government [grant number BCG IT764-13]; MINECO/FEDER, UE [grant number SAF2015-65034-R]; Red de Trastornos Adictivos UE/ERDF [grant numbers RD12/0028/0004 and RD16/0017/0012]; University of the Basque Country [UPV/EHU UFI11/41. NLW held an AI-HS scholarship and Dr. T. Chen Fong scholarship from the Hotchkiss Brain Institute. AWL holds post-doctoral fellowships from AIHS and CIHR. AVH holds Vanier-Canada, AI-HS and BONF studentships. MNH holds a Canada Research Chair Tier 2.

## Materials and Methods

### Animals

All animal care protocols were approved by the University of Calgary’s Animal Care and Use Committee in accordance with the Canadian Council on Animal Care guidelines. Experiments were performed on 21–37 day old male Sprague Dawley rats housed on a 12 hour light/dark cycle with access to Purina Laboratory Chow and water *ad libitum*. Wild type mice (C57BL/6J), conditional pannexin-1 knockout mice ^18^ and TRPV1 knockout mice ^66^ were housed under the same conditions. Panx1 knockout was achieved by tamoxifen delivered daily by peritoneal injections of 100 mg/kg for 5 days ^18^. Controls for the Panx1 knockouts were littermates that received vehicle alone (5% ethanol/95% corn oil) daily for 5 days. Animals were sacrificed at least 72 hours after final injections.

### Chemicals and reagents

All salts used for the artificial cerebral spinal fluid (aCSF) were from Sigma-Aldrich. Capsazepine (10 µM) and anandamide (50 µM), were from Tocris Bioscience. Specific mimetic peptides of Panx1, ^10^panx and a scrambled control peptide of ^10^panx (sc^10^panx) were custom synthesized by AnaSpec or New England Peptide. The C-terminal anti-Panx1 (0.25ng/μl) was from Invitrogen (catalog #488100, rabbit polyclonal). Anti-connexin-43 (0.3ng/μl) was from Abcam (catalog #ab11370, rabbit polyclonal). All drugs were dissolved in water, DMSO or ethanol and aliquoted and frozen. Drugs were then dissolved in aCSF to their final working concentrations. Final concentrations of DMSO or ethanol did not exceed 0.1%.

### Acute hippocampal slice preparation

Rats or mice were anesthetized by isoflurane inhalation in air and decapitated; the brain was extracted, blocked, mounted on a vibrating slicer (VT1200S; Leica) and submerged in an ice-cold high sucrose solution consisting of the following (in mM): 87NaCl, 2.5 KCl, 25 NaHCO3, 0.5 CaCl2, 7 MgCl2, 1.25 NaH2PO4, 25 glucose, and 75 sucrose, saturated with 95% O2 /5% CO2. Transverse hippocampal slices were cut (370 µm for rats and 300 µm for mice) and placed into a chamber containing artificial cerebral spinal fluid at 33°C for at least 1 h before use. aCSF consisted of 120 mM NaCl, 26 mM NaHCO3,3 mM KCl, 1.25 mM NaH2PO4, 1.3 mM MgCl2, 2 mM CaCl2, and 10 mM glucose and was saturated with 95% O2 /5% CO2.

### Primary hippocampal culture preparation

Post natal day zero (P0) pups were used for primary hippocampal cultures as described previously ^17,18^. Pups were anaesthetized on ice and decapitated. Hippocampi were carefully dissected and suspended in a growth media consisting of 1 mM BME supplemented with sodium pyruvate, 50 mg/mL penicillin, 50units/mL streptomycin, B17 supplement (all from Invitrogen), 10 mM HEPES (Sigma), 0.3% glucose (Sigma), and 4% fetal bovine serum. Neural tissue was dissociated by triturating hippocampi in growth media containing papain (Worthington). The neuron / glia co-culture was then washed and plated with growth media on poly-D-lysine and laminin coated 12 mm diameter glass coverslips (Thermo) at 7000–8800 cells/cm^2^.

Cultures were grown in 24-well plates and maintained at 37°C in a humidified incubator (Thermo). Experiments were performed on neuronal cultures from 10 to 20 days after plating, and were equilibrated in aCSF (33°C, bubbled with 95% O_2_ / 5% CO_2_) for at least 30 minutes prior to experimentation.

### Electrophysiology

Slices were transferred to a recording chamber and constantly perfused with aCSF (33°C-35°C) at a rate of 1-2 mL/min. Visualization of hippocampal CA1 pyramidal neurons was achieved with differential interference contrast (DIC) microscopy with an Olympus BX51Wi microscope. Whole cell voltage clamp recordings were performed using borosilicate glass microelectrodes (Sutter Instrument) with a tip resistance of 3-6MΩ which were pulled using a P-1000 Flaming/Brown Micropipette Puller (Sutter Instrument). Microelectrodes were filled with an intracellular solution containing 108 mM potassium gluconate, 2 mM MgCl2, 8 mM sodium gluconate, 8 mM KCl, 2.5 mM K2 -EGTA, 4 mM K2-ATP, and 0.3 mM Na3-GTP at pH 7.25 with 10 mM HEPES. Some experiments were performed with a Panx1 antibody (0.25ng/μl αPanx1) or 50μM anandamide in the pipette. When either of these where included in the intracellular solution the cell was given a minimum of 5 minutes for intracellular equilibration. Electrophysiological data were digitized at 10 kHz and low-pass filtered at 1 kHz with a MultiClamp 700B amplifier and Digidata 1440A analog to digital convertor (Molecular Devices). Data were recorded using pCLAMP 10, Clampex 10.3, and Axoscope 10.3 (Molecular Devices) software and stored for future analysis with Clampfit 10.3, GraphPad Prism 6, and Excel (Microsoft).

Whole-cell current clamp recordings were obtained from CA1 pyramidal neurons held just below spiking threshold (which is approximately -47mV for CA1 pyramidal neurons) by adjusting the holding potential of the neuron with current injections ranging from 50 pA – 120 pA. Membrane potential was recorded for 10 minutes to obtain the baseline, followed by an additional 10 minutes with paired pulse stimulations (two 1 ms stimulations delivered 50 ms apart) delivered to the Schaffer collaterals at one-minute intervals. The frequency of action potentials was quantified with Clampfit 10.3. Data were plotted using Graphpad Prism 6.

### AEA / SR101 transport competition

Experiments measuring flux of AEA through Panx1 were performed by combined patch-clamp electrophysiology and 2-photon laser stimulation microscopy (2PLSM). Cultured primary hippocampal neurons were patched in on-cell mode. The recording pipette and bath solutions consisted of aCSF containing a cocktail of ionotropic channel blockers to isolate Panx1 currents (1μM TTX, 50 µM CdCl, 10 mM TEA, 1 mM 4-AP, 10μM Nifedipine, 100μM picrotoxin, and 20μM MK-801 from Sigma) (18). The recording pipette also contained the Panx1-impermeable FITC-Dextran (50μM, 3,000–5,000 MW) and Panx1-permeable SR101 (100μM, 606.71 MW) fluorescent dyes, with some experiments including 50μM AEA and/or αPanx1. FITC-D and SR101 were excited with a coherent chameleon 2-photon laser tuned to 740 nm. Emission signals were split through a beamsplitter (560 nm) and red/green fluorescence was passed through bandpass filters (585/40 and 525/25 nm, respectively). 2PLSM imaging data were acquired with external photomultiplier tubes and processed with Leica Application Suite Advanced Fluorescence. Changes in fluorescence were calculated as ΔF/F (F-F_o_ / F_o_). If a steady, positive change in FITC-D fluorescence was detected (+5% above baseline for over 20 seconds), it was assumed that the membrane had partially ruptured and the recording was discarded. Images were acquired at 512x512 pixel density at 5s intervals with simultaneous voltage clamp recordings. On-cell patches were held at +60mV (equivalent to -60mV internal membrane potential) and a voltage ramp protocol (equivalent to -80mV to +80mV relative to the inside of the cell, 200 ms) was used to activate Panx1. Charge per sweep was quantified using Clampfit 10.3? and was directly plotted against SR101 influx. The rate of change of fluorescence per unit charge over was calculated as slope, allowing us to normalize for a differential distribution of Panx1 channels present in different experiments.

### Mass Spetrometrical Detection of Anandamide Levels

Hippocampal slices underwent a lipid extraction process as previously described ^67^. In brief, tissue samples were weighed and placed in borosilicate glass culture tubes containing 2 ml of acetonitrile with 5 pmol of [^2^H_8_] AEA for extraction. These samples were homogenized with a glass rod, sonicated for 30 min, incubated overnight at -20°C to precipitate proteins, then centrifuged at 1500 g for 5 min to remove particulates. Supernatants were removed to a new glass culture tube and evaporated to dryness under N_2_ gas, re-suspended in 300 μl of acetonitrile to recapture any lipids adhering to the tube and re-dried again under N_2_ gas. The final lipid extracts were suspended in 200 µl of acetonitrile and stored at -80°C until analysis. AEA contents within lipid extracts were determined using isotope-dilution, liquid chromatography-tandem mass spectrometry (LC-MS/MS) as previously described (Qi et al., 2015).

### Fluorescent anandamide (CAY10455) uptake

HEK293T cells (from ATCC) were transfected with a plasmid encoding rat Panx1 and cultured for 48 hours. Cells were seeded on poly-lysine coated coverslips and grown to confluence. Anandamide uptake was assayed under continuous flow conditions with the fluorescent analog CAY10455 (5μM). Cells were imaged at 1 Hz for 20 minutes with 488 nm excitation and 506-560 nm emission. AEA uptake was calculated as the change in fluorescence over baseline (ΔF/F_0_). Panx1 channels were blocked by pre-incubating cells with ^10^panx1 peptide or a scrambled control (100μM) for 20 minutes prior to CAY10455 perfusion.

### Electron Microscopy

5 TRPV1-WT and 5 TRPV1-KO adult mice of either sex were used in this study. TRPV1-deficient mice (C57BL/6 J background; ^66^) were originally from The Jackson Laboratory (Strain Name: B6.129X1-*Trpv1tm1Jul*/J, Bar Harbor, ME).

Experimental animals were genotyped by polymerase chain reaction (PCR) under standard buffer conditions using the primer pair 5’-CCT GCT CAA CAT GCT CAT TG-3’ and 5’-TCC TCA TGC ACT TCA GGA AA -3’ for the wild-type locus. The primer pair 5’- CAC GAG ACT AGT GAG ACG TG -3’and 5’-TCC TCA TGC ACT TCA GGA AA -3’ was used to detect a fragment in the Neo cassette, specific for the mutant TRPV1 locus. All four primers were used together in the reaction mix (94°C/3 min; 35x[94°C/30 sec, 64°C/1 min, 72°C/1 min]; 1x72°C/2 min; 1x10°C hold).

Homozygous TRPV1-/- and wild-type littermates (TRPV1+/+) from heterozygous breedings were used for experiments. They were deeply anesthetized by intraperitoneal injection of ketamine/xylazine (80/10 mg/kg body weight) and then transcardially perfused at room temperature (RT) with phosphate buffered saline (PBS 0.1 M, pH 7.4) for 20 seconds, followed by the fixative solution made up of 4% formaldehyde (freshly depolymerized from paraformaldehyde), 0.2% picric acid and 0.1% glutaraldehyde in phosphate buffer (PB 0.1 M, pH 7.4) for 10–15 minutes. Brains were then removed from the skull and postfixed in the fixative solution for approximately one week at 4°C. Afterwards, brains were stored at 4°C in 1:10 diluted fixative solution until used.

### Preembedding immunogold method for TRPV1 electron microscopy (EM)

Coronal 50μm- thick hippocampal vibrosections were collected in 0.1 M PB at RT. Then, they were preincubated in a blocking solution of 10% bovine serum albumin (BSA), 0.1% sodium azide and 0.02% saponine prepared in Tris-HCl buffered saline (TBS 1X, pH 7.4) for 30 minutes at RT. Sections were incubated with the primary goat TRPV1 antibody (1:100, VR1 (P-19), sc-1249, Santa Cruz Biotechnology) prepared in the blocking solution with 0.004% saponin, for 2 days at 4ºC. After several washes, tissue sections were incubated with 1.4 nm gold-labeled rabbit antibody to goat IgG (Fab fragment, 1:100, Nanoprobes Inc., Yaphank, NY, USA) prepared in the same solution as the primary antibody for 3 hours at RT. Tissue was washed overnight at 4ºC and postfixed in 1% glutaraldehyde for 10 minutes. After several washes with 1% BSA in TBS, gold particles were silver-intensified with a HQ Silver Kit (Nanoprobes Yaphank, NY, USA) for 12 minutes in the dark. Then, sections were osmicated, dehydrated and embedded in Epon resin 812. Finally, ultrathin sections were collected on nickel mesh grids, stained with lead citrate and examined in a PHILIPS EM208S electron microscope. Tissue preparations were photographed by using a digital camera coupled to the electron microscope. Specificity of the immunostaining was assessed by incubation of the TRPV1 antiserum in TRPV1-KO hippocampal tissue in the same conditions as above.

### Statistical analysis of TRPV1 in the CA1 hippocampus

50μm-thick CA1 hippocampal sections from TRPV1-WT and TRPV1-KO mice (n=5 each) showing good and reproducible silver-intensified gold particles were cut at 80 nm. Electron micrographs (18,000– 28,000X) were taken from grids (2 mm x 1 mm slot) with ultrathin sections showing similar labeling intensity indicating that selected areas were at the same depth. Furthermore, to avoid false negatives, only ultrathin sections in the first 1.5 μm from the surface of the tissue block were examined. Positive labeling was considered if at least one immunoparticle was within approximately 30 nm from the plasmalemma.

TRPV1 metal particles on axon terminals were visualized and counted in randomly taken electron micrographs from both animal types. The number of positive terminals was normalized to the total number of terminals in the images to identify the proportion of TRPV1-positive profiles in TRPV1-WT versus TRPV1-KO. Results were expressed as means of independent data points ± S.E.M. Statistical analyses were performed using GraphPad software 5.0 (GraphPad Software Inc, San Diego, USA).

Statistical analysis of parametric data (i.e. kindling rate) was determined by one-way ANOVA with the post hoc Tukey’s test. Nonparametric data was analyzed with either the Wilcoxon matched-pairs signed rank test (for paired data with direct comparisons) or the Kruskal-Wallis test (with Dunn’s multiple comparisons). For both parametric and nonparametric analyses, significance was set at p≤ 0.05. All results are presented as means ±SEM.

### Data Availability

All relevant data are available from the authors.

**Fig. S1.**
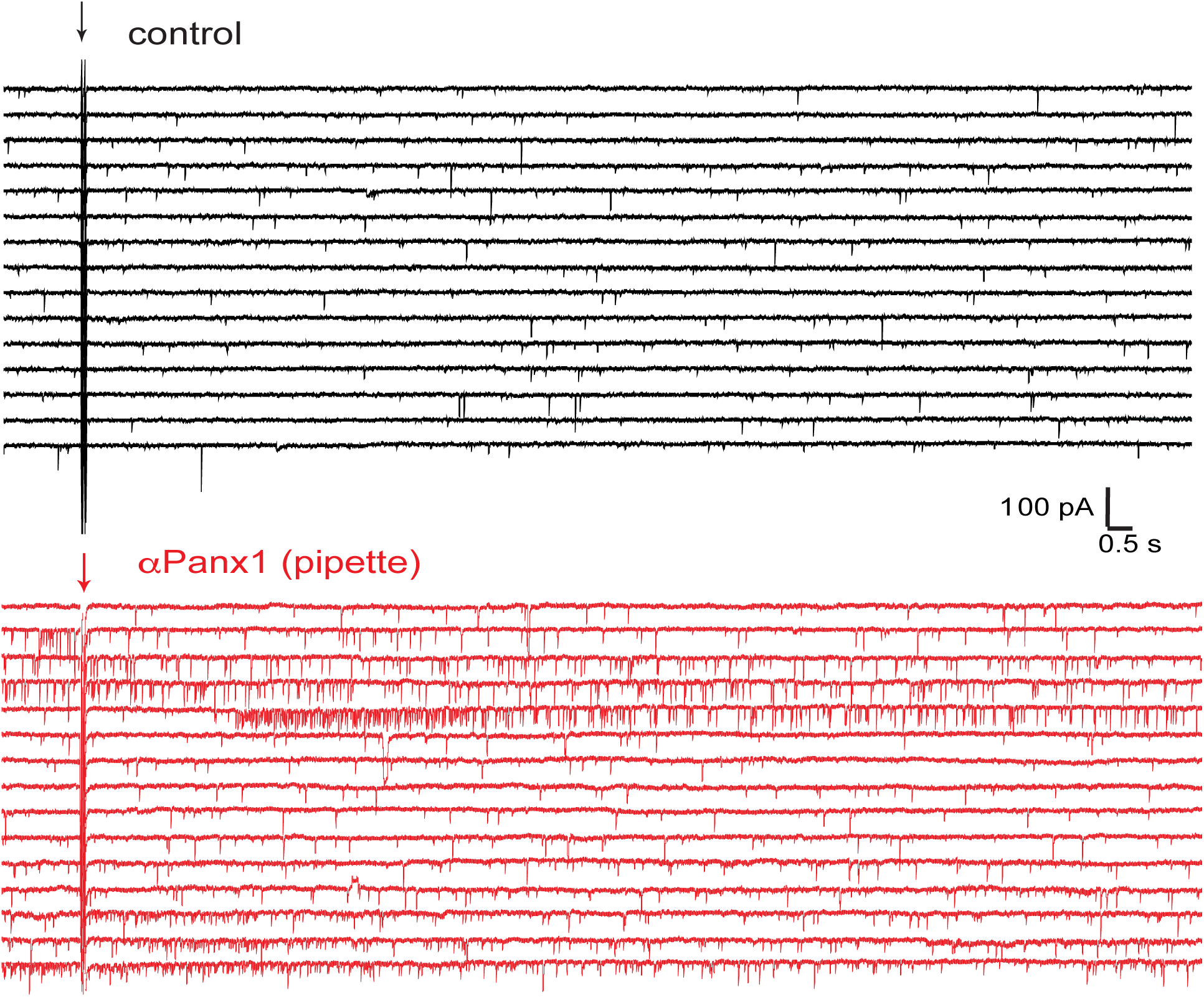
Continuous voltage-clamp recordings from two neurons under control (black) and with the Panx1 blocking antibody, aPanx1 in the pipette (red). Note that when Panx1 was block periods of prolonged synaptic activity are apparent. Picrotoxin is present to block GABAA receptors.

**Fig. S2.**
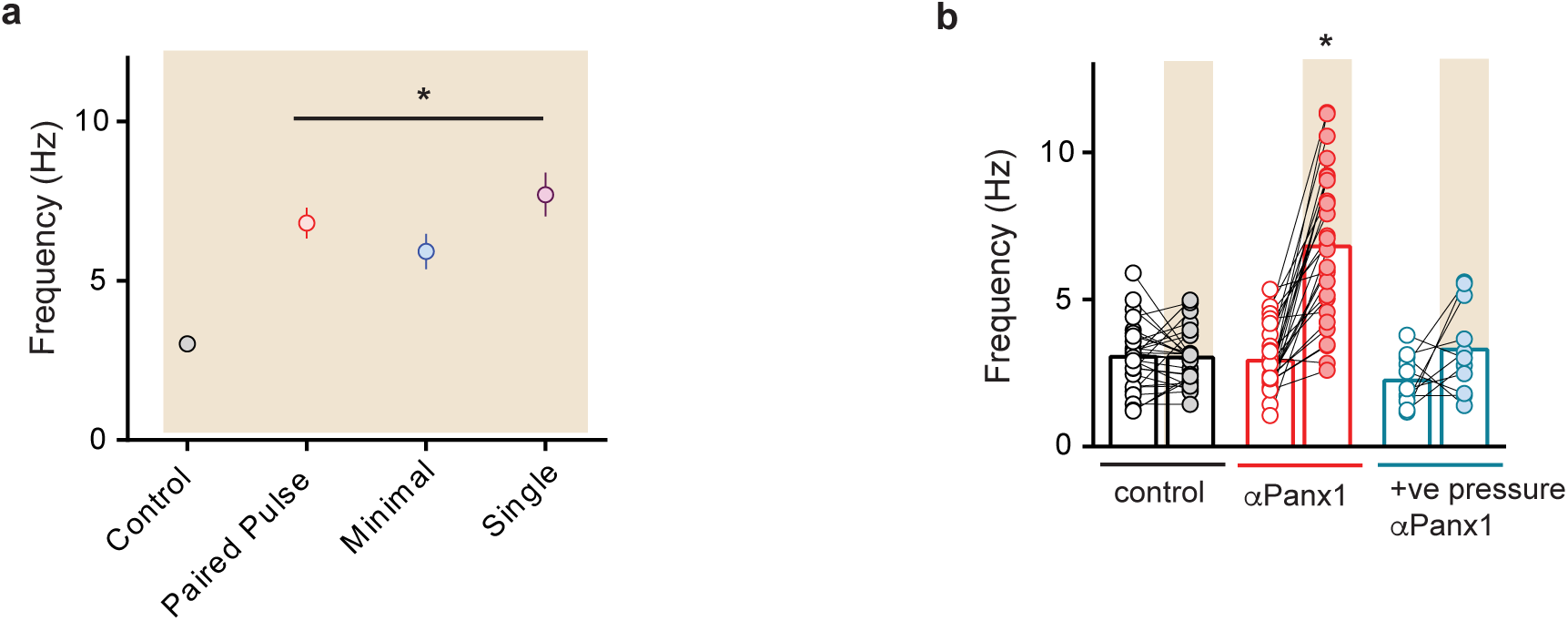
a) Several stimulation paradigms induce asynchronous release when αPanx1 is in the patch pipette. Paired pulse stimulations were 1 ms stims given 50 ms apart every 20 s. Minimal stimulation was 1 ms stims every 20 s with a 50% failure rate. Single stims were 1 ms every 20s at 50% of the maximum eEPSC amplitude. b) comparison of the increase in glutamate neurotransmission when αPanx1 was delivered to the postsynaptic neuron (red) or when it was applied via a patch pipette under positive pressure that was held above the slice. this experiment was to ensure the posiitve pressure did not result in antibody delivery to adjacent neurons to induce prolonged release.

**Fig. S3.**
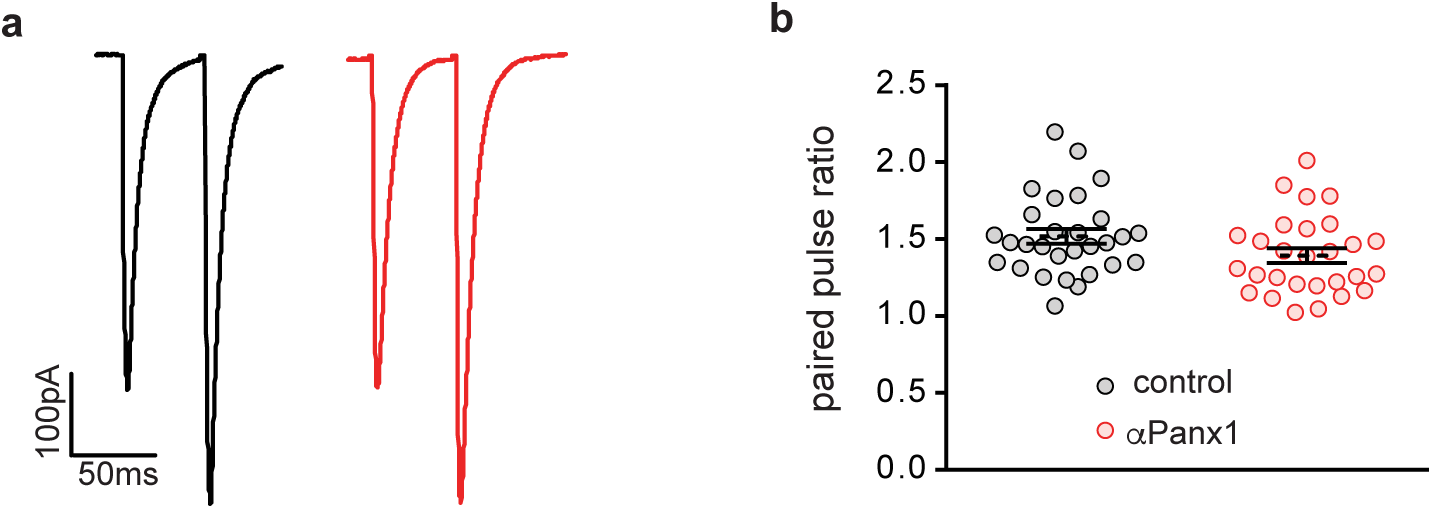
The paired pulse ratio (PPR) was not altered by the presence of postsynaptic αPanx1. a) comparison of paried pulse excitatory postsyanptic potentials under control (black) and with αPanx1 in the pipette (red). b) PPR for all cells tested. Bars are mean +/- sem.

**Figure S4.**
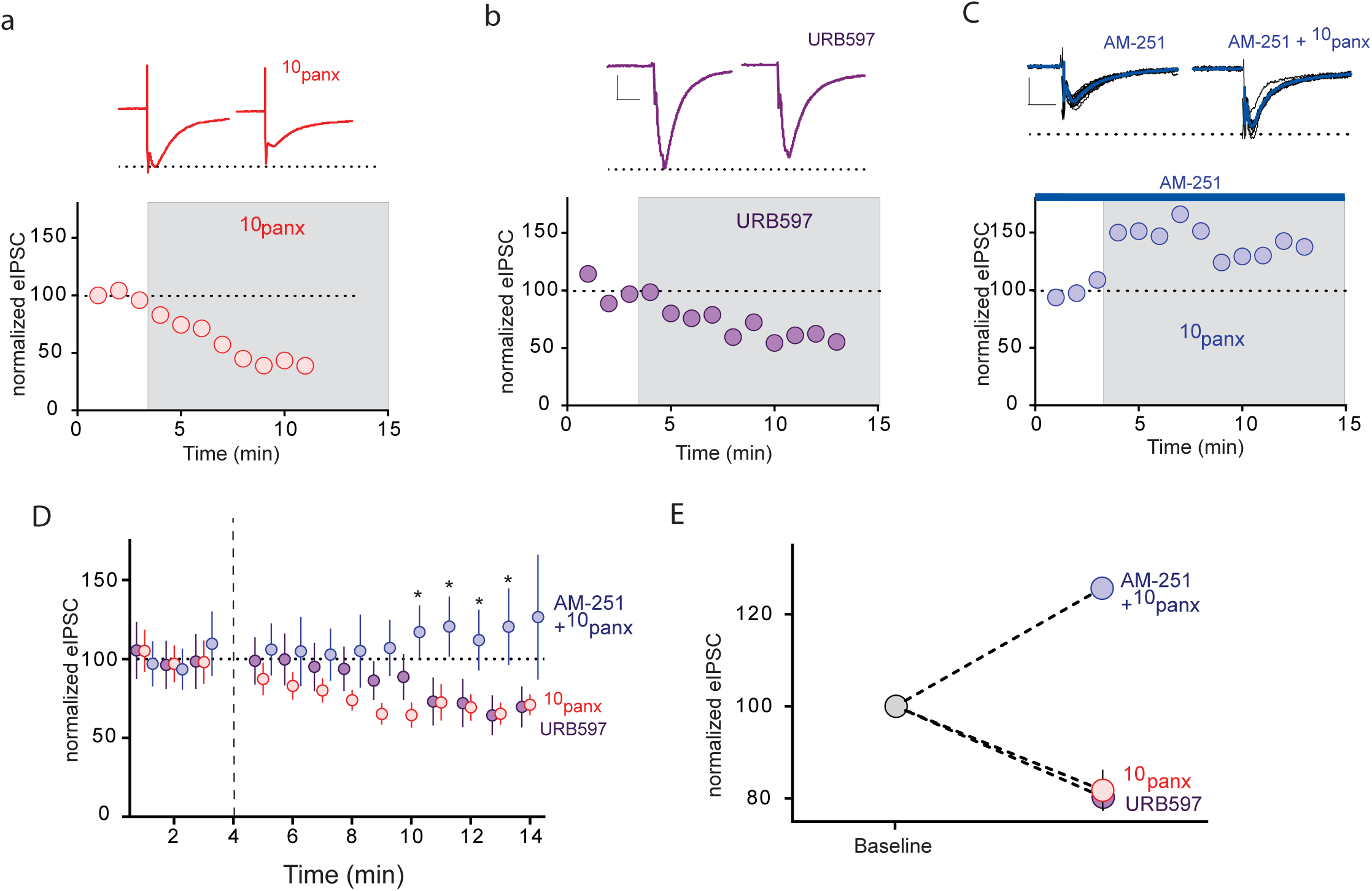
Blocking Panx1 suppresses evoked inhibitory postsynaptic currents that depend upon activation of CB1 receptors. a) Examples of averaged synaptic stimulation induced inhibitory postsynaptic currents (eIPSC) in the absence and presence of the Panx1 blocker, 10panx (100 μM). CA1 neurons were held at -70mV. The time course shows amplitude of eIPSC, averaged at 1 min intervals. The shaded region is when 10panx was applied. b) Examples of averaged eIPSC in the presence of 1 μM URB597 and the time course of inhibition of evoked responses. c) The CB1 R partial agonist, 3 μM AM-251 reversed the inhibition of eIPSCs seen in the presence of 10panx. d) Mean±sem eIPSC amplitudes versus time for each blocker. The gap in the time course occurred during solution switching and equilibration with the drug and no synaptic stimulations were applied. *denotes P<0.05 by 2 way ANOVA. e) Evoked IPSC amplitudes were averaged for the 9–14 min period presented in D. Note that block of FAAH and Panx1 reduced eIPSC amplitudes. Error bars are hidden by the symbols. Scale bars in a-c are 50 pA and 100 ms.

**Fig S5.**
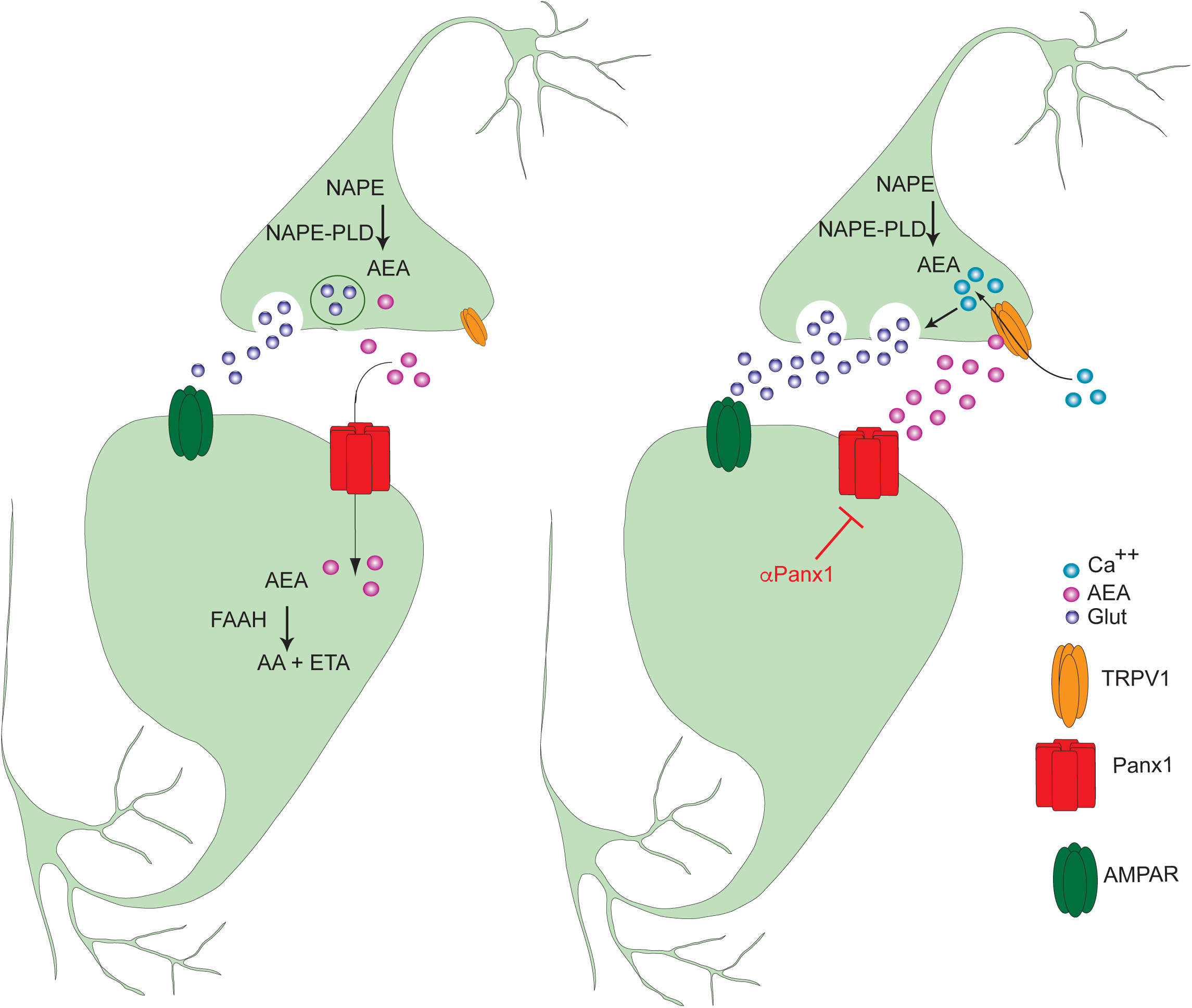
A model of the proposed role of post-synaptic Panx1 and presynaptic TRPV1 in mediating prolonged glutamate release following stimulation. Under basal conditions, AEA is produced in a calcium-dependent way by NAPE-PLD, released and rapidly cleared into postsynaptic neurons via Panx1 for degredation by FAAH. The simplest explanation of our data is that NAPE-PLD is presynaptic, but we cannot completely rule out other glial sources. When Panx1 is blocked, AEA accumulates to activate presynaptic TRPV1 to cause prolonged glutamate release and enhance excitability.

